# Unrestricted eye movements strengthen causal connectivity from hippocampal to oculomotor regions during scene construction

**DOI:** 10.1101/2021.09.23.461606

**Authors:** Natalia Ladyka-Wojcik, Zhong-Xu Liu, Jennifer D. Ryan

**Affiliations:** Department of Psychology, University of Toronto, Toronto ON, CANADA; Department of Behavioral Sciences, University of Michigan-Dearborn, Dearborn MI, USA; Rotman Research Institute, Baycrest Health Sciences, Toronto ON, CANADA; Department of Psychiatry, University of Toronto, Toronto ON, CANADA

**Keywords:** scene construction, eye movements, medial temporal lobes, hippocampus, oculomotor system, dynamic causal modeling

## Abstract

Scene construction is a key component of memory recall, navigation, and future imagining, and relies on the medial temporal lobes (MTL). A parallel body of work suggests that eye movements may enable the imagination and construction of scenes, even in the absence of external visual input. There are vast structural and functional connections between regions of the MTL and those of the oculomotor system. However, the directionality of connections between the MTL and oculomotor control regions, and how it relates to scene construction, has not been studied directly in human neuroimaging. In the current study, we used dynamic causal modeling (DCM) to interrogate effective connectivity between the MTL and oculomotor regions using a scene construction task in which participants’ eye movements were either restricted (fixed-viewing) or unrestricted (free-viewing). By omitting external visual input, and by contrasting free- versus fixed- viewing, the directionality of neural connectivity during scene construction could be determined. As opposed to when eye movements were restricted, allowing free viewing during construction of scenes strengthened top-down connections from the MTL to the frontal eye fields, and to lower-level cortical visual processing regions, suppressed bottom-up connections along the visual stream, and enhanced vividness of the constructed scenes. Taken together, these findings provide novel, non-invasive evidence for the causal architecture between the MTL memory system and oculomotor system associated with constructing vivid mental representations of scenes.

**Highlights:**

- The role of eye movements in mentally constructing scene imagery was investigated
- Restricting eye movements impaired vividness of constructed scene imagery
- Making eye movements strengthened connectivity from memory to oculomotor regions

## 1. Introduction

Our ability to form spatial representations in our mind’s eye is key for supporting navigation, memory, and future thinking (Hassabis & Maguire, 2007; Robin et al., 2016). Functional magnetic resonance imaging (fMRI) studies have demonstrated engagement of the parahippocampal place area (PPA) and hippocampus (HPC) in the encoding of scenes (Epstein & Kanwisher, 1998), as well as in scene construction, the mental generation of coherent spatial contexts in the absence of visual input (Hassabis & Maguire, 2007). Specifically, scene construction entails the imagination and visualization of spatially-coherent scenes in our mind’s eye and is thought to be an important process involved in episodic memory (Lind et al., 2014), imagining the future (Mullally & Maguire, 2014), and spatial navigation (Hassabis & Maguire, 2007). The mental construction of scenes requires both the successful retrieval of relevant spatial and sensory information, and also its integration (Summerfield et al., 2010). Together, the PPA and HPC are thought to support scene construction by binding disparate features and objects into integrated representations (Douglas et al., 2017; Maguire & Mullally, 2013). Significant impairments in the ability to vividly visualize novel scenes in mind have been reported both in individuals with amnesia due to bilateral HPC damage (Mullally et al., 2012, 2014), as well as in individuals with Alzheimer’s disease who have medial temporal lobe (MTL) degeneration (Irish et al., 2015).

Parallel lines of evidence from the field of vision science suggest that saccadic eye movements may play a key role in the construction of scene representations (Kowler, 2011). Contributions of the oculomotor system to scene construction have received limited investigation (see Mirza et al., 2016; Parr & Friston, 2017), largely because prior neuroimaging studies have typically instructed participants to close their eyes (Hassabis et al., 2007; Mullally et al., 2012); consequently, standard eyetracking techniques could not be used to measure eye movements that were made while imagining scenes. Saccade motor maps accurately code for locations of objects in space (Zimmermann & Lappe, 2016). Adaptively changing the targeting position of saccades (i.e., changing the required saccade amplitude) subsequently disrupts localization of objects in visual space, suggesting that scene perception and memory may rely on an oculomotor map (Bahcall & Kowler, 2000; Ryan & Shen, 2020). Conversely, saccades and corresponding gaze fixations are guided by prior knowledge regarding the expected locations of objects within scenes (Castelhano & Heaven, 2011). Patterns of gaze fixations during imagination are similar to those during perception, suggesting that oculomotor behavior supports mental imagery (Gurtner et al., 2021) by reinstating previously encoded spatiotemporal content (Wynn et al., 2019). Individuals move their eyes across a blank screen in accordance with object positions during recall of previously studied scenes (Johansson et al., 2006) and when merely listening to auditory scene descriptions (Spivey et al., 2000). Eye movements therefore support construction of mental representations even in the absence of external visual input (Conti & Irish, 2021).

Recent fMRI findings highlight the functional connectivity between the PPA and early visual regions (Baldassano et al., 2013). Computational modeling has revealed vast structural connections between the MTL and oculomotor control regions, including the frontal eye fields (FEF) (Ryan et al., 2020a; Shen et al., 2016). These connections are functionally relevant; simulated stimulation of HPC subfields and parahippocampus resulted in rapid evoked responses in the FEF (Ryan et al., 2020a). In humans, single-pulse transcranial magnetic stimulation of FEF created top-down activity that directly shaped responses in lower-level visual regions (Veniero et al., 2021). However, to our knowledge, no study has investigated the direction of information flow among the MTL, FEF, and early visual regions in human neuroimaging, and specifically, during scene construction. Particularly, empirical evidence is needed to demonstrate that brain regions that support mental representations or the construction of mental representations can have causal, top-down influence on the oculomotor control system and ultimately affect early visual regions. Testing this directional information flow is crucial for us to understand not only how we mentally construct scenes, but also how we navigate in a familiar environment, imagine future scenarios, or recall past spatial or contextual information.

The present study used combined eyetracking-fMRI recordings and manipulations of viewing behavior to elucidate interactions among the MTL, FEF, and visual cortex during scene construction. Participants were prompted with word labels of scenes and instructed to freely move their eyes around a blank screen (free-viewing) or to maintain fixation (fixed-viewing) while imagining the cued scene. Since there was no strong visual input in this task (e.g., compared to scene viewing tasks), the design allowed us to examine more clearly the directionality of information flow among MTL, FEF, and early visual regions when internal representations are used to construct scenes. Specifically, dynamic causal modeling (Friston et al., 2003) was used to interrogate directionality of effective couplings (i.e., effective connectivity) between regions and how the directional coupling can be modulated by the viewing manipulation.

In contrast to functional connectivity, which can only reveal the degree to which component regions correlate over time, DCM can be used to test how these regions causally interact with one another. DCM refers to the inversion of generative (i.e., forward) models of observable brain responses (e.g., fMRI signals) that combine a biologically plausible neural model and established hemodynamic models of fMRI signals. Briefly, first, in the so-called bilinear neural model, the neural activity change rate at any moment in each brain region is modeled and determined by the summed effects of (1) the baseline directional influence from other connected regions in the model, i.e., effective connectivity effects, (2) the rate of neural activity self-decay within individual regions due to self-inhibition, (3) the change in the effective connectivity effects due to experimental manipulation (in the current experiment, from fixed- to free-viewing), and (4) direct external stimulation if applied to this region (for detailed explanations, see section 2.8. Dynamic Causal Modeling analysis and Stephan & Friston, 2010; Zeidman et al., 2019a).The neural model is then combined with a biologically plausible hemodynamic model to produce fMRI BOLD signals. Finally, the observed fMRI signal from all regions will be used to inverse this combined dynamic forward model and produce parameter estimates for effective (i.e., causal or directional) connectivity among the regions of interest (ROIs) and the effective connectivity changes due to experimental condition manipulation, as well as self-inhibition within each region and other parameters involved. Because this is a generative model and parameters are defined based on models with clear biological meanings, DCM allows us to test how different regions causally affect each other during baseline and how this effective connectivity changes in different experimental conditions.

Because prior work has shown that neural activity in the PPA and HPC (Liu et al., 2020) scaled with increasing gaze fixations, whereas restricting fixations reduced neural activity (Liu et al., 2017, 2020), by comparing the free-viewing versus fixed-viewing conditions, we could then assess, in the present study, the changes in the strength of the directionality of information flow between regions. This allowed us to understand how the MTL interacts with oculomotor control regions such as FEF when mental representations are used to construct scenes. Compared to fixed-viewing, free-viewing during scene construction was hypothesized to strengthen top-down connections from the PPA and HPC towards the FEF and lower-level visual regions. When viewing was restricted (fixed-viewing), we predicted suppression of bottom-up connections from early visual regions towards oculomotor regions. This work highlights the interaction of the MTL and oculomotor system in the active construction of scenes.

## 2. Material and methods

### 2.1. Participants

Thirty-three healthy young adults (18 female) aged 18 to 30 (age: *M* = 22.97 years, *SD* = 3.31; education: *M* = 16.29 years, *SD* = 1.93) from the University of Toronto and surrounding Toronto area community completed this experiment in exchange for monetary compensation. The sample size was determined based on our previous investigations on the relationship between eye movement and brain activity (Liu et al., 2017, 2020; Wynn et al., 2021) and the literature on experimental fMRI and DCM methods (Goulden et al., 2012; Murphy & Garavan, 2004; Szucs & Ioannidis, 2020). Thirty-one subjects had participated in a scene viewing task earlier in the same scanning session, as reported in Liu et al. (2020). All participants had normal or corrected-to-normal vision (including color vision), and none had any neurological or psychological conditions. The study was approved by the Research Ethics Board at Rotman Research Institute at Baycrest Health Sciences.

### 2.2. Stimuli

Stimuli consisted of 28 unique word labels for common semantic scene categories (e.g., casino, ski resort, etc.). These word labels were presented in the center of the screen followed by either a green fixation dot (free-viewing condition) or a red fixation dot (fixed-viewing condition) (Figure 1).

**Figure 1:**
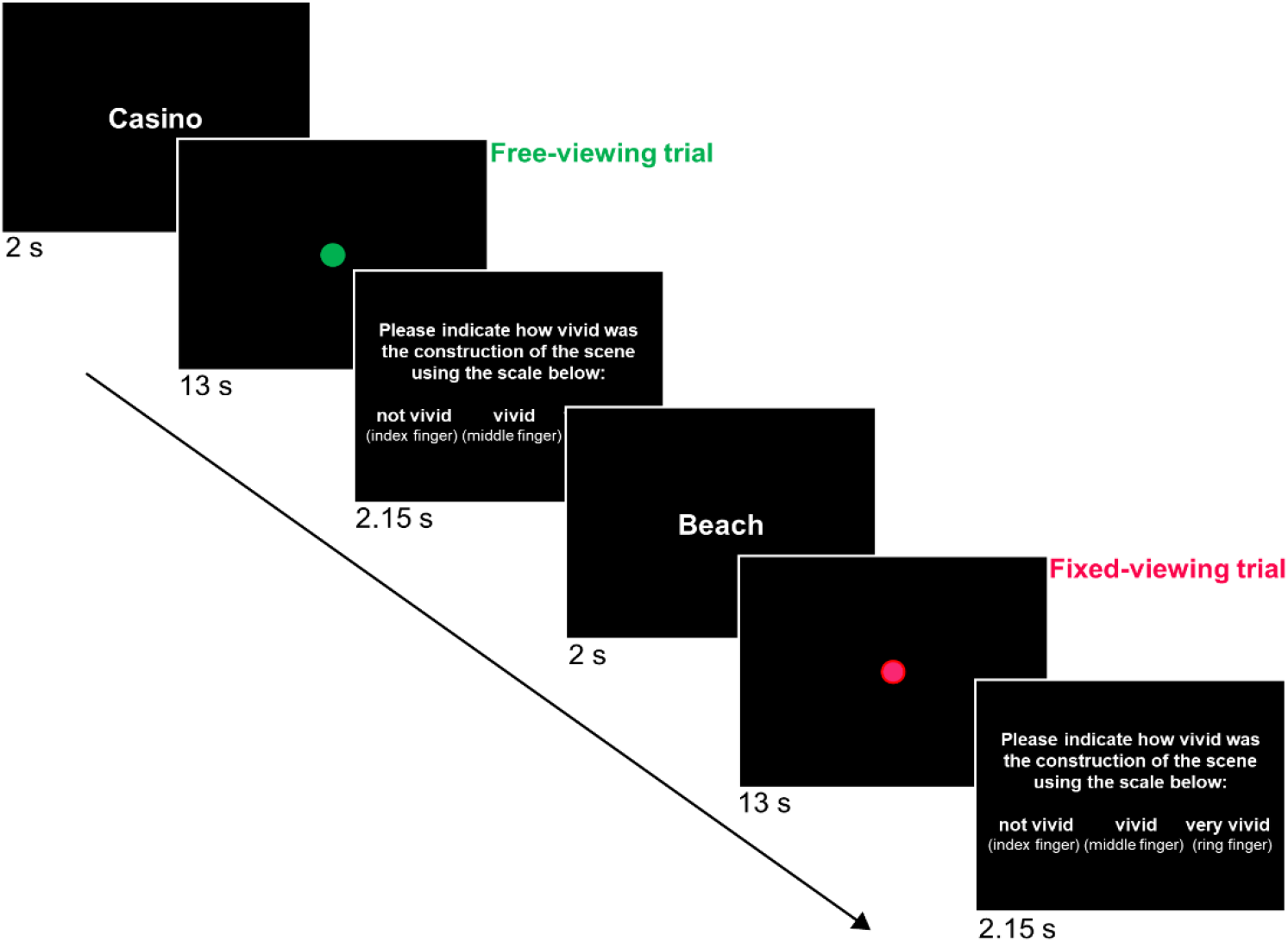
Scene construction task procedure. Participants were presented with a word cue of a scene and then instructed to either freely move their eye gaze across the screen (free-viewing) or to keep their eye gaze fixed on the fixation dot (fixed-viewing), as they mentally constructed the cued scene. Following the scene construction period, participants responded to the vividness of their mental construction with a 3-button response for not vivid, vivid, or very vivid.

### 2.3. Procedure

Participants completed one run containing 28 trials in the scanner. Half of the trials were studied under free-viewing instructions and half were studied under restricted (i.e., fixed) viewing instructions, in a randomized order (Figure 1). At the start of each trial, participants were shown, for 2 seconds, a word label of a scene category that they were to mentally construct. The word labels were counterbalanced over the two viewing conditions across participants. Next, a fixation dot was presented at the center of the screen for 13 seconds. The color of the fixation dot indicated the viewing condition for the trial; a green fixation dot indicated a free-viewing trial and a red fixation dot indicated a fixed-viewing trial. In the free-viewing condition, participants were instructed to freely explore the blank screen as they wished for the duration of the trial as they were mentally constructing a scene based on the word label cue. In the fixed-viewing condition, participants were instead required to keep their eye gaze at the location of the fixation dot while mentally constructing the cued scene. Following each scene construction trial, participants were given 2.15 seconds to respond to a vividness rating question using an MRI-compatible button box. Possible vividness ratings were: 1 (not vivid), 2 (vivid), and 3 (very vivid).

All stimuli were presented with Experiment Builder (Eyelink 1000; SR Research) back-projected to a screen (projector resolution: 1024×768) and viewed with a mirror mounted on the head coil.

### 2.4. Eyetracking

During the scene construction task, monocular eye movements were recorded inside the scanner using the EyeLink 1000 MRI-compatible remote eyetracker with a 1000Hz sampling rate (SR Research Ltd., Mississauga, Ontario, Canada). The eyetracker was placed inside the scanner bore (behind the participant’s head) and detected the right pupil and corneal reflection via a mirror mounted on the head coil. To ensure successful tracking during the task, a nine-point calibration was performed at the beginning of the scanning session and online manual drift correction was performed between trials when necessary. EyeLink’s default eye movement event parser was used to categorize fixations and saccades. A velocity threshold of 30°/s and an acceleration threshold of 8000°/s were used to classify saccades (saccade onset threshold = 0.15°). Events not defined as saccades or blinks were classified as fixations. The number of fixations that participants made during scene construction was calculated and exported to a MATLAB-compatible environment using the EyeLink software Data Viewer for further analyses.

### 2.5. MRI scan acquisition

As specified in Liu et al. (2020), a 3T Siemens MRI scanner with a standard 32-channel head coil was used to acquire structural and functional MRI images. T1-weighted high-resolution MRI images for structural scans were obtained using a standard 3D MPRAGE (magnetization-prepared rapid acquisition gradient echo) pulse sequence (176 slices, FOV = 256 × 256 mm, 256×256 matrix, 1 mm isotropic resolution, TE/TR = 2.22/2000 ms, flip angle = 9 degrees, and scan time = 280 s). For the functional scan, BOLD signal was assessed using a T2*-weighted EPI acquisition protocol with TR = 2000 ms, TE = 27 ms, flip angle = 70 degrees, and FOV = 192 × 192 with a 64 × 64 matrix (3 mm x 3 mm in-place resolution; slice thickness = 3.5 mm with no gap). A total of 250 volumes were acquired for the fMRI run, with the first 5 discarded to allow the magnetization to stabilize to a steady state. Both structural and functional images were acquired in an oblique orientation 30° clockwise to the anterior–posterior commissure axis.

### 2.6. fMRI data preprocessing

The fMRI preprocessing procedure was previously reported in Liu et al. (2020) and is reproduced here. SPM12 (Statistical Parametric Mapping, Wellcome Trust Centre for Neuroimaging, University College London, UK) in the MATLAB environment (The MathWorks Inc., Natick, USA) was used to process the functional images. Following standard SPM12 preprocessing procedure, slice timing was first corrected using *sinc* interpolation with the reference slice set to the midpoint slice. Next, functional images were aligned using a linear transformation, and for each participant functional image parameters from the alignment procedure (along with global signal intensity) were checked manually using the toolbox ART (http://www.nitrc.org/projects/artifact_detect/). Anatomical images were co-registered to the aligned functional image and segmented into white matter, gray matter, cerebrospinal fluid, skull, and soft tissues using SPM12’s default 6-tissue probability maps. Segmented images were then used to calculate the transformation parameters mapping from the subjects’ native space to the MNI template space. The resulting transformation parameters were used to transform all functional and structural images to the MNI template. The functional images were finally resampled at 2×2×2 mm resolution and smoothed using a Gaussian kernel with an FWHM of 6 mm. The first five fMRI volumes from each run were discarded to allow the magnetization to stabilize to a steady state.

### 2.7. GLM fMRI analysis

We used SPM12 to conduct the first-level (i.e., individual) whole brain General Linear Model (GLM) analysis, comparing brain activation differences between the free-viewing and fixed-viewing conditions. We separately convolved trials with duration = 13 seconds in the free-viewing and fixed-viewing condition with the canonical hemodynamic function (HRF) in SPM12, which served to be the 2 main regressors of interest. We added 6 motion parameters obtained from the co-registration process as regressors of no interest. Default high-pass filters with a cutoff of 128 s were applied and serial correlations were removed using a first-order autoregressive model AR(1). Next, to examine differences in neural responses that were elicited by gaze fixations, we contrasted the free-viewing with the fixed-viewing condition for each participant. Then, at the group-level, individual participants’ above-described contrast estimates were entered into one-sample *t* tests.

For this analysis, we first focused on two *a priori* ROIs, i.e., HPC and PPA (see Figure 2). Based on our previous findings (Liu et al., 2017; 2020), we hypothesized that both regions should show stronger activity in the free-versus fixed-viewing condition. For the HPC mask, FreeSurfer’s *recon-all* function (https://surfer.nmr.mgh.harvard.edu/; Fischl, 2012) was used to extract subject-specific anatomical masks for all 33 participants. For the PPA, group-level masks were defined functionally based on an earlier scene-scrambled picture processing task (i.e., scene versus scrambled images) reported in Liu et al. (2020). Note that for the PPA, this group-level mask was determined from 31 of the total 33 participants who had previously participated in a picture processing task in Liu et al. (2020) during the same testing session. The MNI coordinates for the peak activation in the right PPA were [32, −34, −18] and peak activation in the left PPA were [-24, −46, −12]. The left and right PPA mask contained 293 and 454 (1×1×1 mm^3^) voxels, respectively. In this analysis, for each participant, the mean beta estimates were extracted within each ROI for each contrast and then we used one-tailed *t* tests to test our *a priori* hypotheses at the group-level.

**Figure 2:**
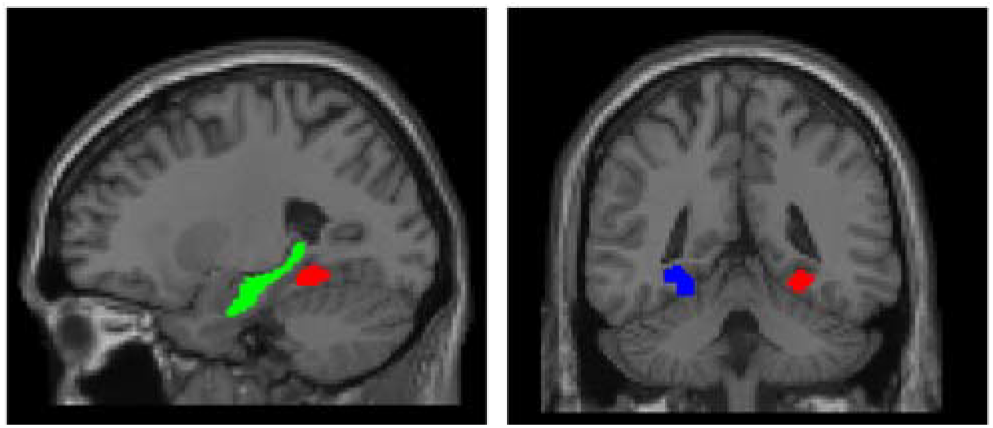
ROI masks used in the present study. Masks are shown in MNI space (RPI orientation), with the left HPC in green*, left PPA in red, and right PPA in blue. *For illustrative purposes, a single subject’s mask is displayed for the HPC.

For completeness, we also examined voxel-wise whole brain results to explore regions showing different engagement in the two conditions (i.e., *free-viewing* – *fixed-viewing*). The statistical threshold was set to *p* = .005 with 10 voxel extension (uncorrected) to facilitate future meta-analyses (Lieberman & Cunningham, 2009). The automated anatomical labeling (AAL) toolbox (Tzourio-Mazoyer et al., 2002) was used to identify anatomical labels for regions that showed significant effects.

### 2.8. Dynamic Causal Modeling analysis

#### 2.8.1. Model design

In the present study, we used dynamic causal modeling (DCM) to assess the directionality of information flow among regions of the MTL involved in scene representations and regions involved in oculomotor control, along with early visual regions. DCM for fMRI has been shown to have high scan-rescan reliability (Schuyler et al., 2010) and lends itself to modeling neural activity separately from BOLD responses (Stephan & Friston, 2010). Since DCM estimates region-specific hemodynamic responses and neural states, it is also less susceptible to issues related to HRF variations in different brain regions (Friston et al., 2014; Stephan et al., 2007). Moreover, compared to other measures of effective connectivity, such as structural equation modeling (Büchel & Friston, 1997; McIntosh & Gonzalez-Lima, 1994) or Granger causality (Goebel et al., 2003), DCM presents an advantage in that it allows for the direct evaluation of modulatory effects of contextual (i.e., task) conditions. Here, we focused on three separate models to investigate the top-down influences of the MTL and the bottom-up influences of early visual regions on the oculomotor system.

First, we focused on a 3-ROI model with the PPA, FEF, and primary visual cortex (V1). This model (see Figure 3A) was selected to investigate how top-down signals from the scene processing region PPA towards the oculomotor control region FEF may be modulated when participants were allowed to freely move their eyes. Specifically, we hypothesized that allowing free eye movements during scene construction would strengthen the directional connections from the PPA to the FEF, and that this would, in turn, drive the activation of V1, typically the first stage of cortical processing of visual information.

**Figure 3:**
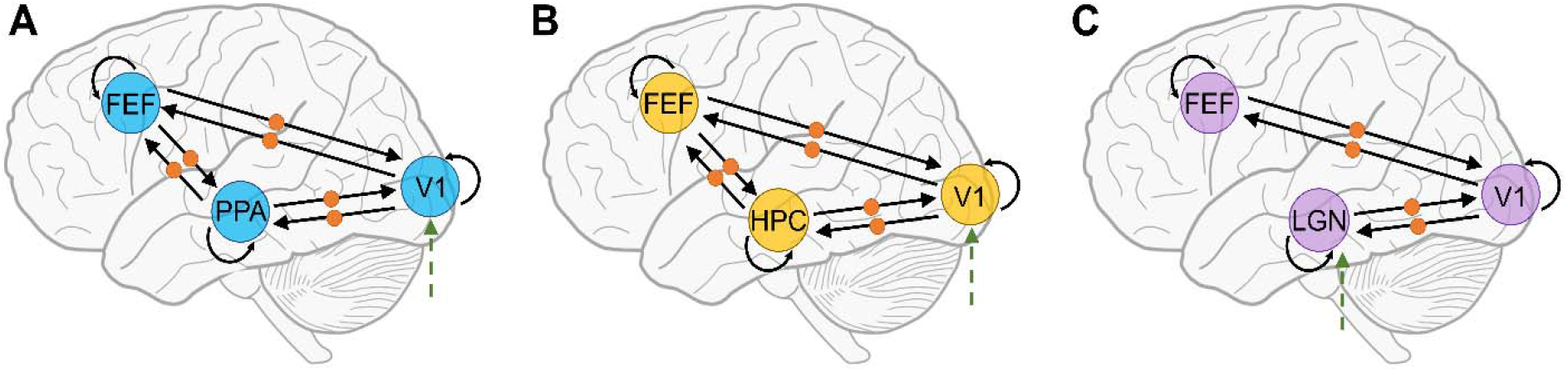
Full dynamic causal model (DCM) spaces. (A) Full model for the PPA, FEF, and V1 ROIs. (B) Full model for the HPC, FEF, and V1 ROIs. (C) Full model for the LGN, V1, and FEF ROIs. For each model, the solid straight arrows represent baseline connections between regions and the curved solid arrows represent intrinsic self-connections (A matrix in the state equation of DCM). The orange circles on the arrows for each model represent modulatory connections (B matrix in the state equation of DCM) and the dotted green arrows represent the driving input (C matrix in the state equation of DCM).

Next, to examine how the hippocampus specifically interacts with the oculomotor control and early visual regions, we specified a 3-ROI model with the HPC, FEF, and V1 (see Figure 3B). Like the above-described model with the PPA, we were interested in investigating how topdown signals from the memory representation region HPC towards the FEF may be modulated by free-viewing versus fixed-viewing during scene construction. As for the PPA, we hypothesized that allowing free viewing would strengthen the top-down connections from the HPC to the FEF, and from the HPC and FEF to V1. Because there is already robust evidence showing that the PPA and HPC closely interact with each other, in this study, we did not put the two regions in the same model to avoid complex models and to focus on our main questions of interest.

In the third model, we explored whether the top-down modulation effect arising from free-versus fixed-viewing would be evident in the earliest visual processing pathway region in the central nervous system, i.e., the lateral geniculate nucleus (LGN) in the thalamus. Therefore, we constructed a 3-ROI model with the LGN, V1, and FEF (see Figure 3C). We excluded the connection between the FEF and LGN based on low prior evidence in humans and nonhuman primates for connectivity between these two regions (Gilbert & Li, 2013; Kashihara, 2020).

Due to the increased complexity in interpretation when including inter-hemispheric connections for DCM (Stephan et al., 2010; Stephan & Friston, 2010; Zeidman, Jafarian, Seghier, et al., 2019), we designed our models for the right and left hemispheres separately.

#### 2.8.2. Time series extraction for DCM

We used the significant group-level activation clusters to extract time series for the DCM analysis, which were constrained to be within the boundaries of our ROIs. For the PPA and HPC, we used the same masks as in the GLM ROI analyses to constrain boundaries. For all other ROIs, we used peak activation coordinates in the activated clusters in these regions (i.e., FEF, V1, and LGN) from the group-level GLM results, which we subsequently confirmed using Neurosynth (Yarkoni et al., 2011). Specifically, the location of the FEF was based on both the anatomical landmark (i.e., the superior frontal sulcus intersection with the precentral gyrus; Vernet et al., 2014) and MNI coordinates found in the literature ([L: −22 −10 50; R: 20 −9 49]; Donner et al., 2000). Both V1 and LGN were located based on their anatomical landmarks on the MNI template. Next, an 8-mm radius sphere centered at peak activation was used for each participant to mark the ROI location, then voxels within that sphere that showed free- vs. fixed-viewing effect (*p* = 0.05, no corrections) were used as the final ROIs. Including voxels that show task modulation effects can facilitate DCM analysis (Zeidman, Jafarian, Corbin, et al., 2019). However, when the number of voxels was lower than 50 (i.e., too few voxels in the ROI) for a specific ROI of a specific participant, we relaxed the threshold to 0.1, 0.5, or without using any threshold (i.e., threshold = 1) until at least 50 voxels could be obtained for that ROI and that participant. The same threshold procedure was applied to the HPC and PPA to ensure a sufficient number of voxels included in the DCM analysis.

After ROIs were defined, SPM12 was used to extract BOLD signals at each voxel of an ROI and then the first principal component of the time series data from all voxels in the ROI was computed for the DCM analysis.

#### 2.8.3. Individual-level DCM analysis

DCM implemented in SPM12 was used for effective connectivity analysis. DCM for fMRI models the dynamics of the neural states underlying the BOLD response by a differential state equation that describes how these responses change with the current neural states, contextual conditions, and driving inputs (Friston et al., 2003). The advantage of DCM is that it not only provides a measure of endogenous effective connectivity between regions, but also a measure of how these directed connections are modulated by task demands (Stephan & Friston, 2010). The state equation for DCM is:

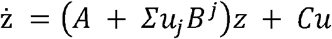

The intrinsic and baseline interactions between the neuronal states are endogenous connections and quantified by A parameters. These interactions are mediated by anatomical connections and are irrespective of task condition (i.e., when the two conditions are combined). The influences of the task conditions on connectivity between ROIs are modulations and quantified by B parameters, i.e., connections altered on free-viewing trials over fixed-viewing trials. The influences of driving inputs are quantified by C parameters (in this case, caused by all trials). In this study, we used a mean-centered driving input so that the matrix A parameters represented an average effective connectivity across experimental conditions and matrix B modulatory parameters add or subtract from this average (i.e., strengthening or weakening the average, or baseline, connectivity). By using this mean-centered approach, we were able to focus on how the free- and fixed-viewing conditions modulate the directionality of functional underlying couplings.

As shown in Figure 3, the PPA and HPC models are configured to have bidirectional baseline connections between connected ROIs (A matrix), which is consistent with the literature on the anatomical and functional connectivity between these regions. Within-ROI auto-connections in the A matrix were also added by default (Zeidman, Jafarian, Corbin, et al., 2019). Since we were interested in how the eye movement conditions modulate effective connectivity between regions, we included all possible modulatory connections between the combinations of ROIs (B matrix) for the free-viewing condition. For all of these model designs, the driving inputs (C matrix) were set to enter the earliest ROI in the visual stream (i.e., V1 in the PPA – FEF – V1 and HPC – FEF – V1 models, and LGN in LGN – V1 – FEF model).

#### 2.8.4. Group-level DCM analysis

In line with previous work using DCM (Zeidman, Jafarian, Seghier, et al., 2019), we used a Parametric Empirical Bayes (PEB) approach to evaluate group effects (i.e., commonality among participants) on connectivity parameters. With different modulatory parameters switched on or off, the model space for PPA – FEF – V1 and HPC – FEF – V1 consisted of 64 possible models, and the model space for LGN – V1 – FEF had 16 possible models. This PEB process was implemented using Bayesian Model Reduction and then averaging the parameters from the best reduced models with Bayesian Model Averaging. Specifically, the winning model was selected on the basis of offering the best fit to the data with the highest exceedance probability, which denotes the probability that this model is more likely than any other in the given dataset. This analysis produced weighted model parameters for the winning model, which we report along with connectivity matrices. Since this approach relies on simultaneous estimation of nested models with Bayesian inference, we did not need to correct for multiple comparisons (Gelman & Tuerlinckx, 2000; Stephan et al., 2007). As suggested by Kass and Raftery (1995), we reported model parameters with posterior probabilities (Pp) above 95% which corresponds to strong evidence in favour of a model.

## 3. Results

### 3.1. Eye movements during fMRI scanning

To confirm the effect of the eye movement manipulation, we compared the average number of gaze fixations (Figure 4A) and the average saccade amplitude (Figure 4B) that participants made in the free-viewing and fixed-viewing conditions. Paired *t*-tests revealed that participants made a greater number of gaze fixations (*t*(32) = 7.01, *p* < .001, *d* = 1.22) in the free-viewing than the fixed-viewing condition, and that participants had a larger saccade amplitude measured in degrees of visual angle (*t*(32) = 7.00, *p* < .001, *d* = 1.24) in the free-viewing than the fixed-viewing condition. The fixation frequency across participants for the free-vs. fixed-viewing conditions based on the location within the blank screen (1024×768 pixels) is displayed in Figure 4D.

**Figure 4:**
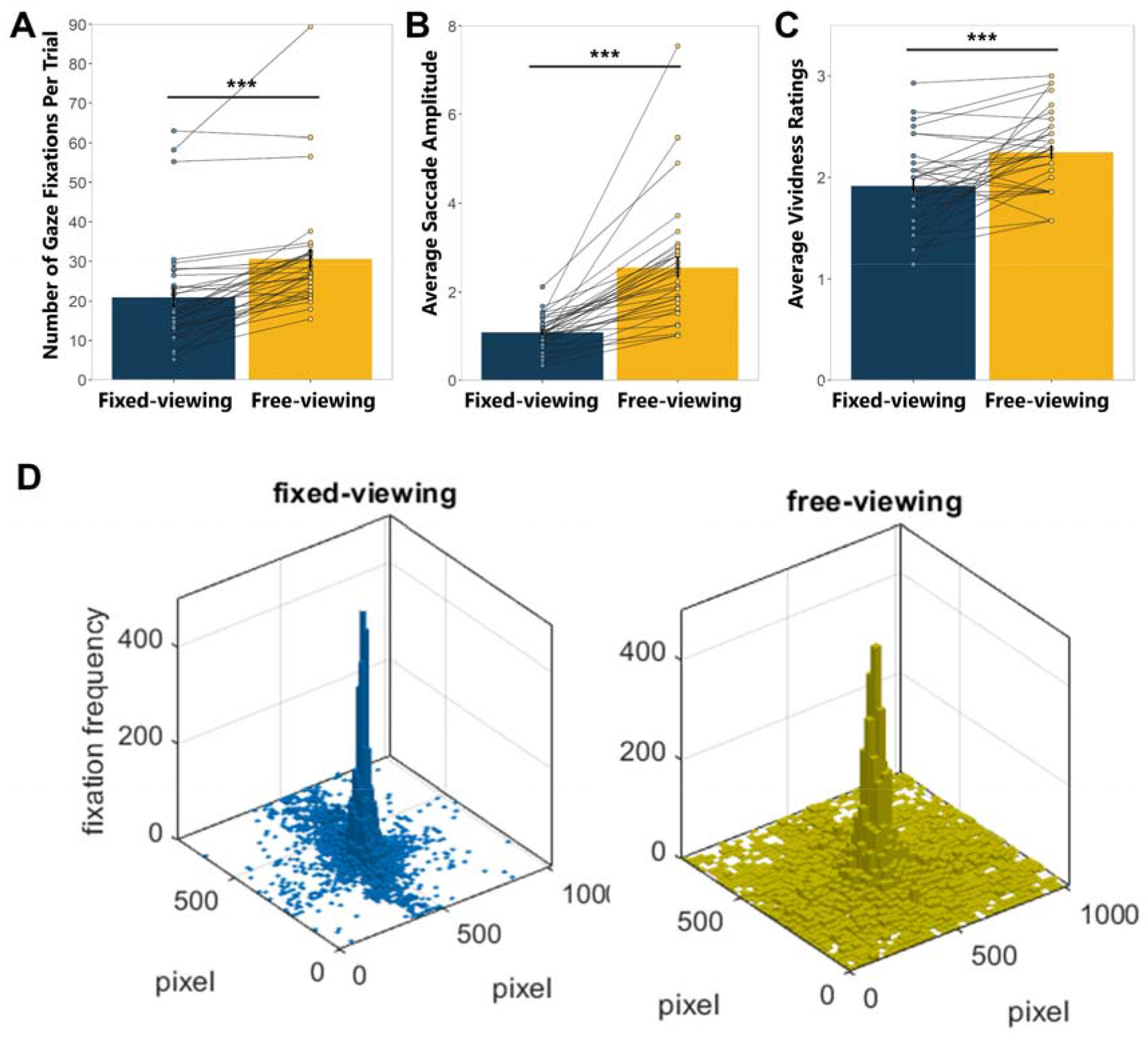
Eye-tracking results for free- vs. fixed-viewing. (A) Average number of gaze fixations per trial in the fixed-viewing and free-viewing conditions. More fixations were elicited in the free-viewing condition. Lines represent each subject’s averaged fixations across trials. (B) Average saccade amplitude in the fixed- and free-viewing conditions. Greater saccade amplitude was elicited during the free-viewing condition. Lines represent each subject’s averaged saccade amplitude across trials. (C) Average vividness rating in the fixed- and free-viewing conditions. Constructed scenes were rated more vivid during the free-viewing condition. Lines represent each subject’s averaged vividness ratings across trials. (D) Fixation frequency based on location within the blank screen (1024×768 pixels) during fixed-viewing (left) and free-viewing (right) trials across participations. Error bars are ± SEM. \*\*\**p* <.001

### 3.2. Vividness Ratings

At the end of each trial of the scene construction task, participants were asked to provide a vividness rating ranging across 1 (not vivid), 2 (vivid), and 3 (very vivid). We compared the average vividness rating for the fixed- and free-viewing conditions. A paired *t*-test revealed that participants rated vividness for trials in the free-viewing condition (*M* = 2.25, *SD* = 0.39) higher than for trials in the fixed-viewing condition (*M* = 1.92, *SD* = 0.42; *t*(32) = 4.13, *p* < .001, *d* = 0.72; Figure 4C).

### 3.3. GLM fMRI Results

We first examined the brain activation contrast between free- and fixed-viewing in the PPA and HPC to confirm whether activity in our *a priori* ROIs of the PPA and HPC were stronger in the free- versus fixed-viewing condition during scene construction. As hypothesized, the left and right PPA showed stronger activation in the free-viewing compared to the fixed-viewing condition (left: *t*(32) = 4.65, *p* < .0001, *d* = 0.81; right: *t*(32) = 3.62, *p* < .001, *d* = 0.63; one-tailed). Additionally, the left HPC showed stronger activation in the free-viewing compared to the fixed-viewing condition (*t*(32) = 1.70, *p* < .05, *d* = 0.30; one-tailed). Our ROI analysis of the right HPC did not yield significant effects (*t*(32) = 0.54, *p* > 0.05; one-tailed; Figure 5A).

**Figure 5:**
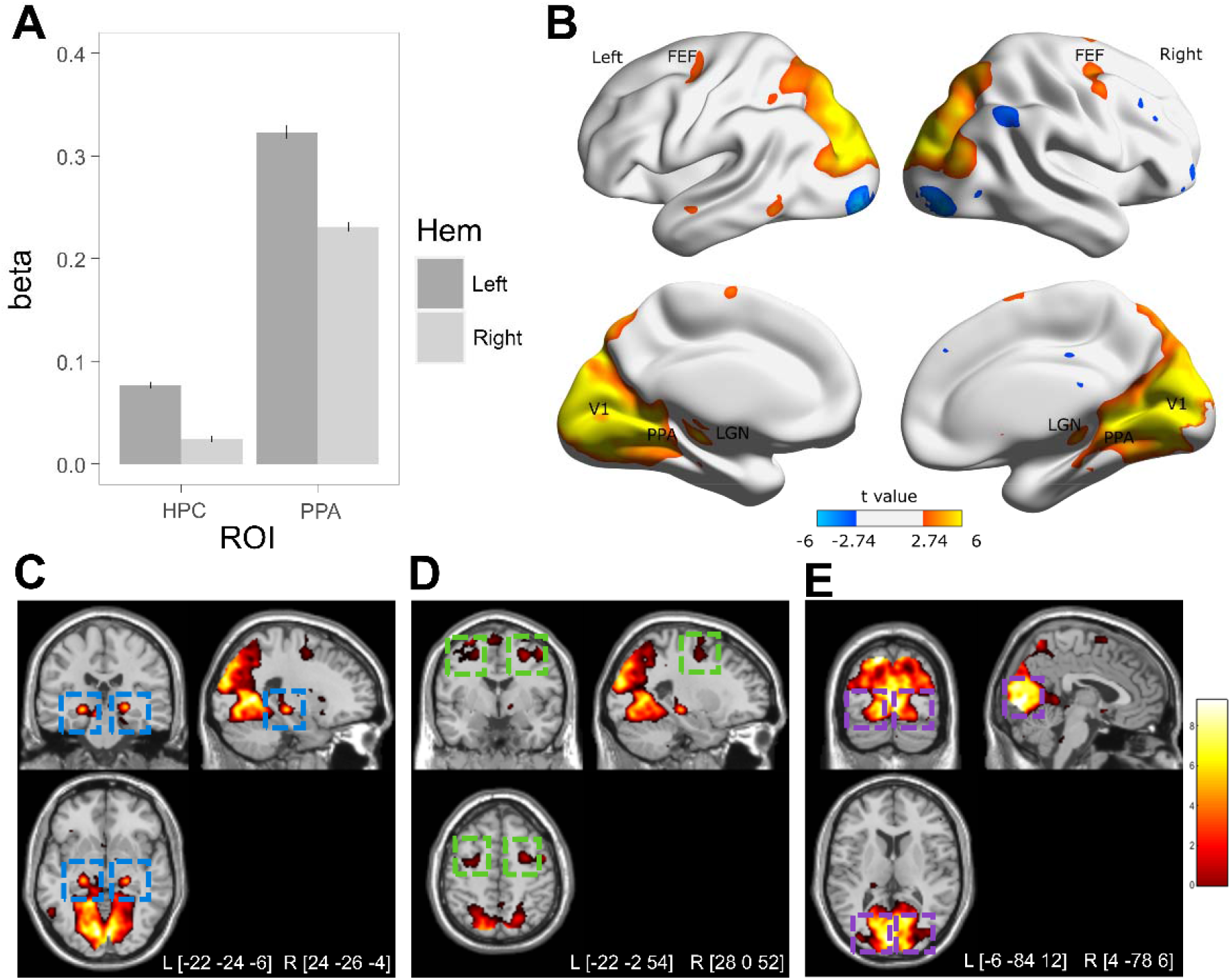
GLM fMRI results. (A) ROI analysis revealed a significant effect of free-viewing over fixed-viewing in bilateral PPA (left *p* <.0001; right *p* <.001; one-tailed), left HPC (*p* <.05; one-tailed), but not in right HPC (*p* > 0.05; one-tailed). Error bars are ± SEM. (B) Brain surface plots showing voxel-wise activation differences between the free-viewing and fixed-viewing conditions. (C) Voxel-wise whole brain results showing stronger activation in the left and right LGN in the free-vs. fixed-viewing condition (highlighted in blue boxes; *ps* < .00001). (D) Voxel-wise whole brain results showing stronger activation in the left and right FEF in the free- vs. fixed-viewing condition (highlighted in green boxes; *p* = .001 and .0003). (E) Voxel-wise whole brain results showing stronger activation in the left and right V1 in the free-vs. fixed-viewing condition (highlighted in purple boxes; *ps* < .00001). For (B), (C), (D), and (E), brain images are thresholded at *p* < .005, 10 voxel extension (no corrections) for illustration purposes. Clusters showed free-viewing > fixed-viewing effects at FEF, PPA, HPC, V1 and LGN are indicated.

The results of the voxel-wise whole brain analysis that showed increased and decreased neural activity during free-viewing compared to fixed-viewing are illustrated in Figure 5B and listed in Table 1 below. The HPC ROI did not survive the whole-brain analysis. As can be seen in Figure 5C, 5D, and 5E, both the ventral and dorsal visual processing pathway regions, including in the bilateral LGN (Figure 5C), bilateral FEF (Figure 5D), and bilateral V1 (Figure 5E), showed stronger activation when participants were allowed to freely move their eyes.

**Table 1.**
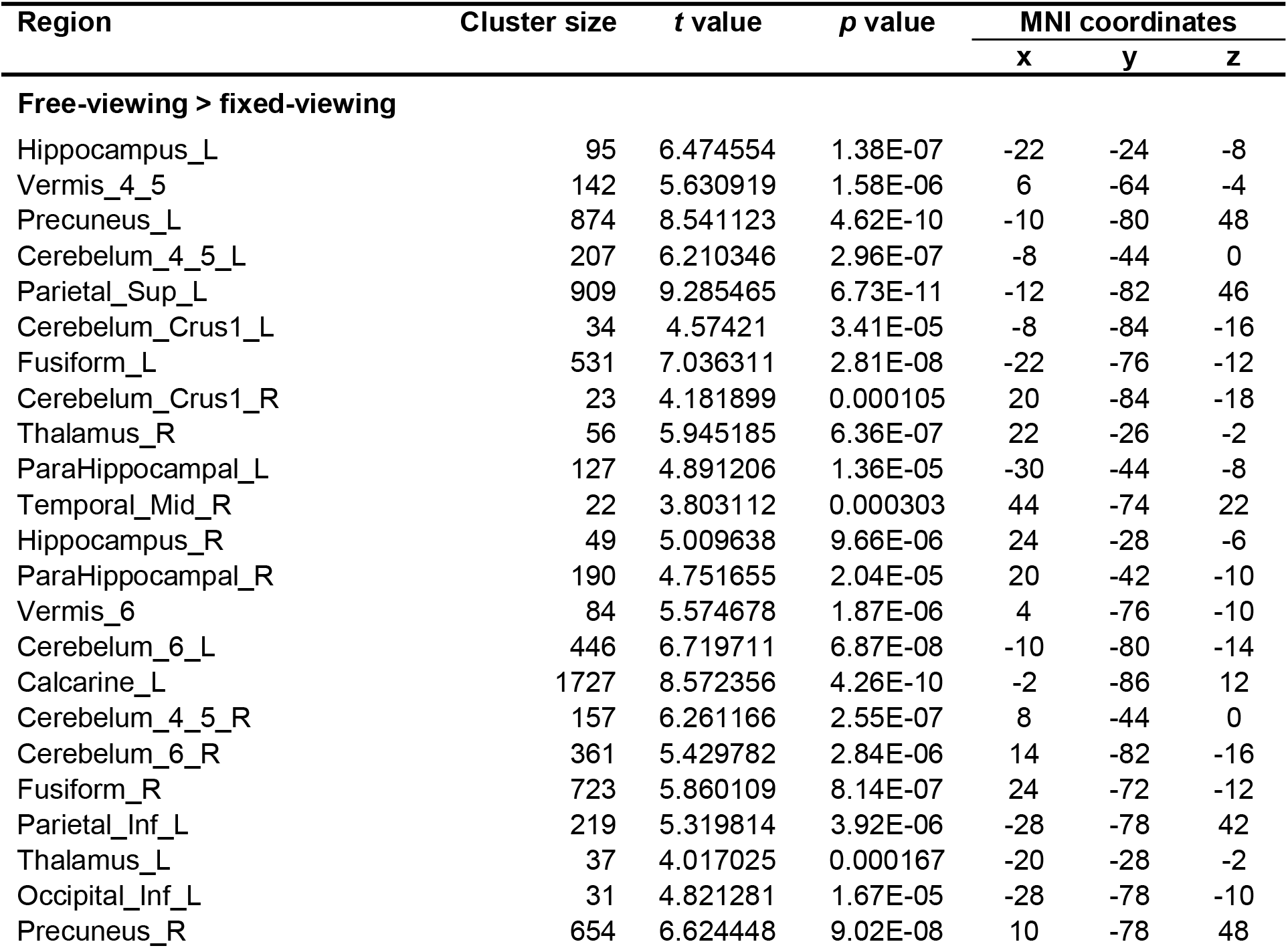
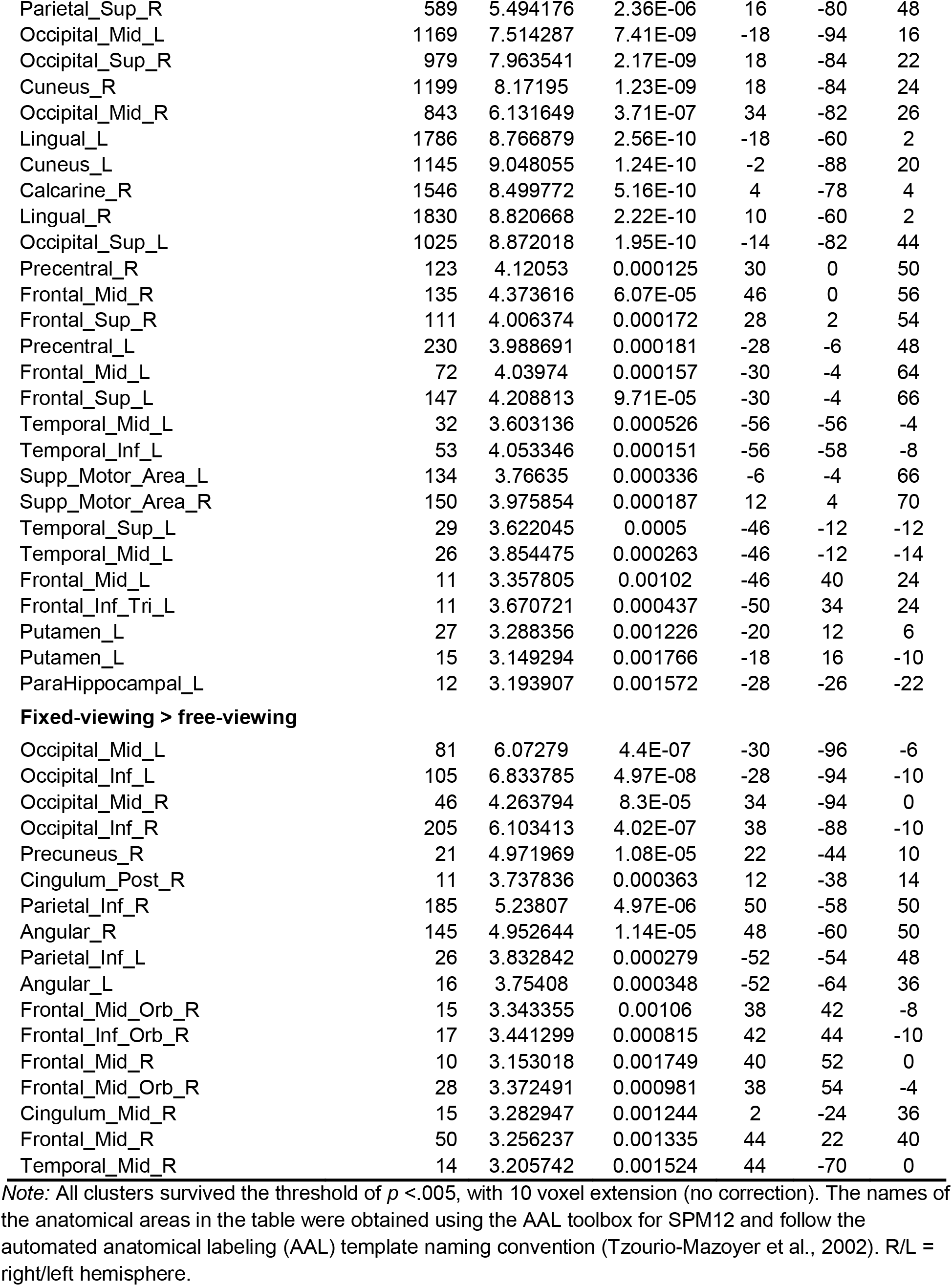
Brain regions that showed stronger and weaker activation in the free-viewing than fixed-viewing condition.

### 3.4. DCM Results

#### 3.4.1. Model 1: PPA – FEF – V1

As shown in Figures 6A and 6C (i.e., the A matrix parameter results), our DCM analyses found bidirectional excitatory baseline connectivity between the scene processing region PPA and early visual region V1, and between the oculomotor control region FEF and V1, during scene construction when the two viewing conditions were averaged. There was also an excitatory connection directed from FEF to PPA (in the left hemisphere) and an inhibitory connection from PPA to FEF (in the right hemisphere), but no excitatory effect from PPA to FEF. However, when participants were allowed to freely move their eyes during scene construction (i.e., the free-viewing condition), the PPA showed enhanced excitatory connectivity to FEF (*B*_PPA→FEF_ = 0.876 and 0.582 for the left and right hemisphere, *Pp* > 95%; see green arrows in Figure 6B and 6D). Furthermore, both PPA and FEF showed enhanced excitatory effects on the early visual region V1, especially in the left hemisphere (*B*_PPA→V1_ = 1.039 and *B*_FEF→V1_ = 0.840, *Pp* > 95%). Taken together, these excitatory connections indicate an enhanced top-down information flow from the scene processing region PPA to the oculomotor control region FEF, and to the early visual region V1. In the right hemisphere, the connectivity from FEF to PPA was also enhanced when participants were allowed to freely move their eyes (*B*_FEF→PPA_ = 0.410, *Pp* > 95%), whereas the bottom-up effect (i.e., the directional connectivity from V1 to PPA and FEF) was weakened during the free-versus fixed-viewing condition (*B*_V1→PPA_ = −0.426/−0.610, *B*_V1→FEF_ = −0.904/−0.758 for the left/right hemisphere, *Pp* > 95%; see red arrows in Figure 6B and 6D).

**Figure 6:**
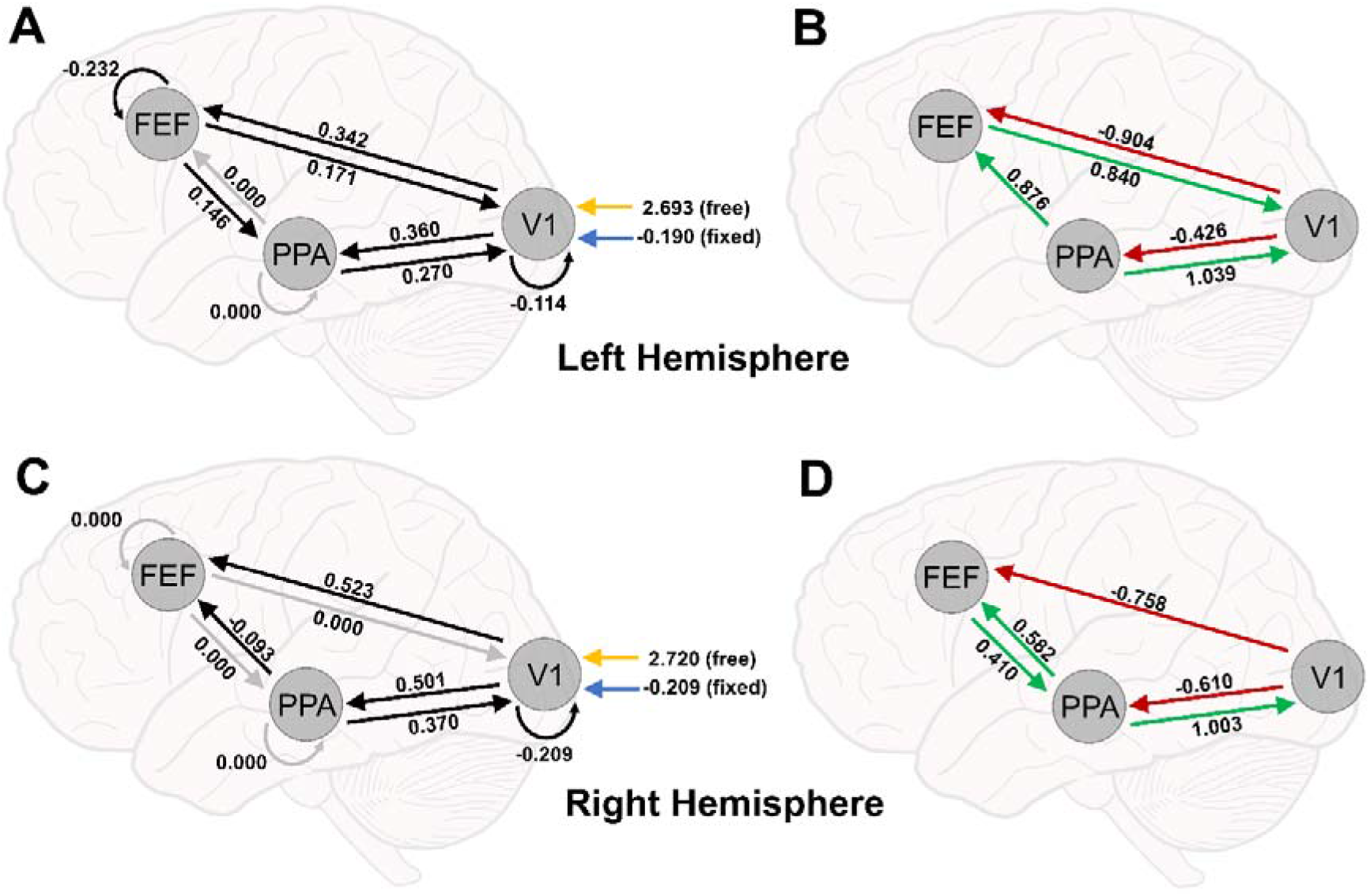
DCM results for PPA – FEF – V1. (A) Baseline connections for the PPA – FEF – V1 model in the left hemisphere. (B) Modulatory connections for the PPA – FEF – V1 model in the left hemisphere. (C) Baseline connections for the PPA – FEF – V1 model in the right hemisphere. (D) Modulatory connections for the PPA – FEF – V1 model in the right hemisphere. For (A) and (C) solid black arrows with a straight line indicate connections that exceeded posterior probabilities (Pp) of 95%, while grey arrows indicate connections that did not exceed a 95% Pp. Straight arrows represent baseline connections between ROIs, and curved arrows represent the strength of self-connections within ROIs, with smaller (i.e., more negative) values indicating less self-inhibition within the ROI (i.e., long activation sustainment). The model inputs (*C* matrix) are indicated as “free” and “fixed” to V1. For (B) and (D) red arrows indicate significant negative modulatory effects and green arrows indicate positive modulatory effects (Pp >95%).

#### 3.4.2. Model 2: HPC – FEF – V1

As shown in Figures 7A and 7C (i.e., the *A* matrix parameter results), we found bidirectional underlying excitatory connectivity in both hemispheres between the memory region HPC and early visual region V1, and between the oculomotor control region FEF and V1, during scene construction when the two viewing conditions were averaged. In the right hemisphere, the DCM analysis found an excitatory baseline connection from HPC to FEF, and an inhibitory connection from FEF to HPC. However, in the left hemisphere, there was no excitatory effect between the HPC and FEF (Figure 7A). When free-viewing was contrasted with the fixed-viewing condition, the HPC showed enhanced excitatory connectivity to FEF in both hemispheres (*B*_HPC→FEF_ = 0.646 and 0.627 for the left and right hemisphere, *Pp* > 95%; Figure 7B and 7D). Additionally, in the left hemisphere, we found an enhanced excitatory effect from FEF to HPC (*B*_FEF→HPC_ = 0.490, *Pp* > 95%). Although there were no enhanced excitatory effects from HPC towards V1 bilaterally, in the right hemisphere we found an enhanced excitatory connection from FEF to V1 (*B*_FEF→V1_ = 0.735, *Pp* > 95%).

**Figure 7:**
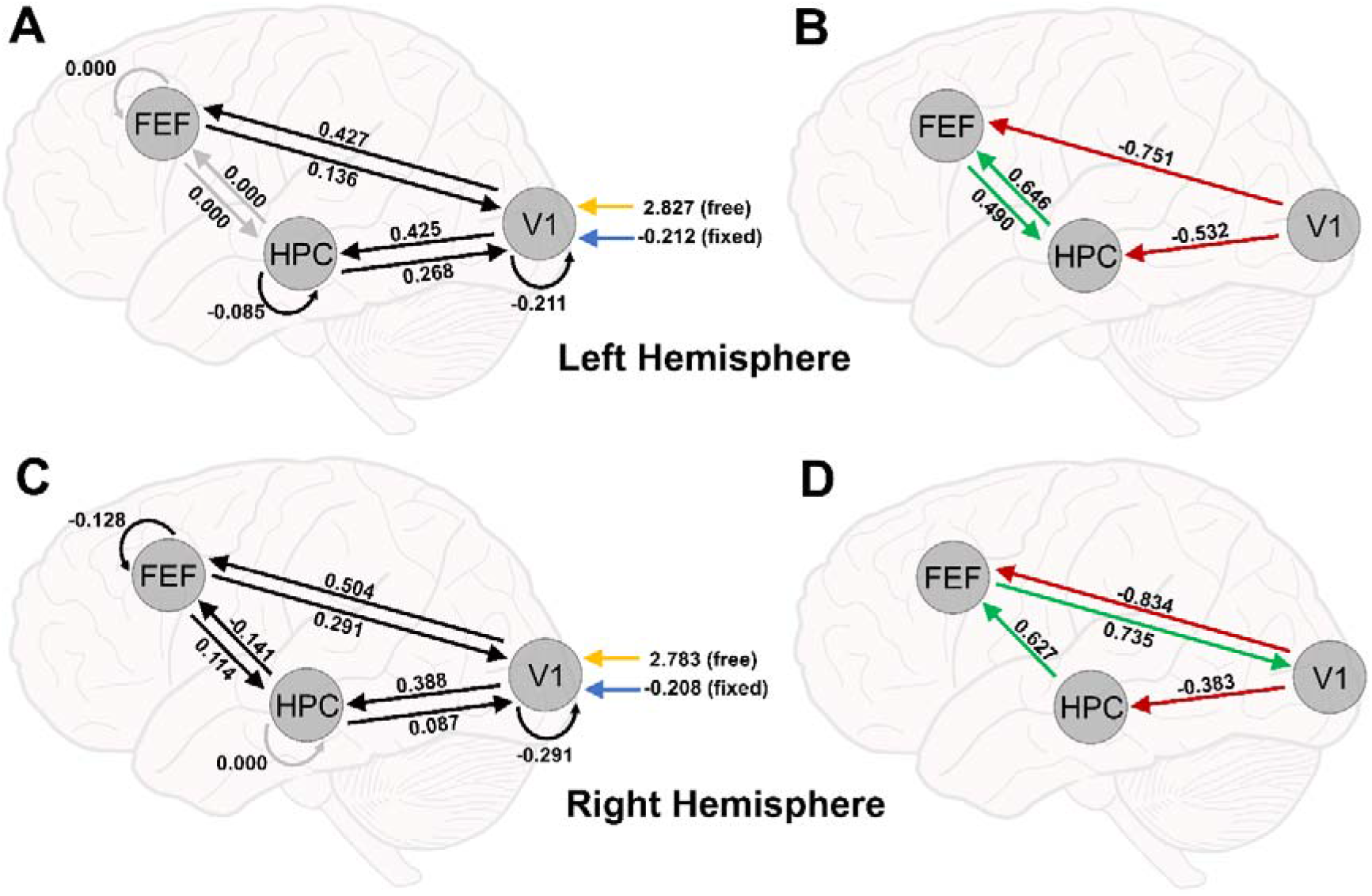
DCM results for HPC – FEF – V1. (A) Baseline connections for the HPC – FEF – V1 model in the left hemisphere. (B) Modulatory connections for the HPC – FEF – V1 model in the left hemisphere. (C) Baseline connections for the HPC – FEF – V1 model in the right hemisphere. (D) Modulatory connections for the HPC – FEF – V1 model in the right hemisphere. For (A) and (C) solid black arrows with a straight line indicate connections that exceeded posterior probabilities (Pp) of 95%, while grey arrows indicate connections that did not exceed a 95% Pp. Straight arrows represent baseline connections between ROIs, and curved arrows represent the strength of self-connections within ROIs, with smaller (i.e., more negative) values indicating less self-inhibition within the ROI (i.e., long activation sustainment). The model inputs (C matrix) are indicated as “free” and “fixed” to V1. For (B) and (D) red arrows indicate significant negative modulatory effects and green arrows indicate positive modulatory effects (Pp >95%).

These excitatory connections indicate an enhanced top-down information flow from the memory region HPC to the oculomotor control region FEF, with enhanced information also flowing from FEF back to HPC, during free-viewing. Finally, the bottom-up effect from V1 to HPC and FEF was weakened during the free-versus fixed-viewing condition (*B*_V1→HPC_ = −0.532/−0.383, *B*_V1→FEF_ = −0.751/−0.834 for the left/right hemisphere, *Pp* > 95%; see red arrows in Figure 7B and 7D), consistent with the results found in the PPA – FEF – V1 model.

#### 3.4.3. Model 3: LGN – V1 – FEF

When the two viewing conditions were averaged during scene construction, we found bidirectional baseline connectivity between V1 and LGN in both hemispheres (Figure 8A and 8C, i.e., the A matrix parameter results). Here, the connection from LGN to V1 was excitatory, whereas the connection from V1 to LGN was inhibitory. Additionally, we found an inhibitory baseline connection from FEF to V1 in both hemispheres, as well as an excitatory connection from V1 to FEF in the left hemisphere. When free-viewing was contrasted with the fixed-viewing condition, the bottom-up influence from the LGN, to V1, and to FEF was weakened (*B*_LGN→V1_ = −1.309/−1.380, *B*_V1→FEF_ = −1.001/−1.176 for the left/right hemisphere, *Pp* > 95%; see red arrows in Figure 8B and 8D). In both hemispheres, the top-down influence from the FEF to V1 was strengthened (*B*_FEF→V1_ = 1.057 and 1.136 for the left and right hemisphere, *Pp* > 95%), as found in the previous models. Interestingly, the modulation effect from V1 to LGN was inhibitory (*B*_V1→LGN_ = −0.777 and −0.816 for the left and right hemisphere, *Pp* > 95%), and the input (mainly from the fixed-viewing trials) had a negative influence on LGN (the blue upward arrow in Figure 8A and 8C, i.e., the C matrix). Taken together, these results indicate that when participants could freely move their eyes during scene construction, the bottom-up information flow was inhibited, and although the enhanced top-down influence occurred, it stopped at the earliest cortical region V1, i.e., did not extend to the thalamic region LGN.

**Figure 8:**
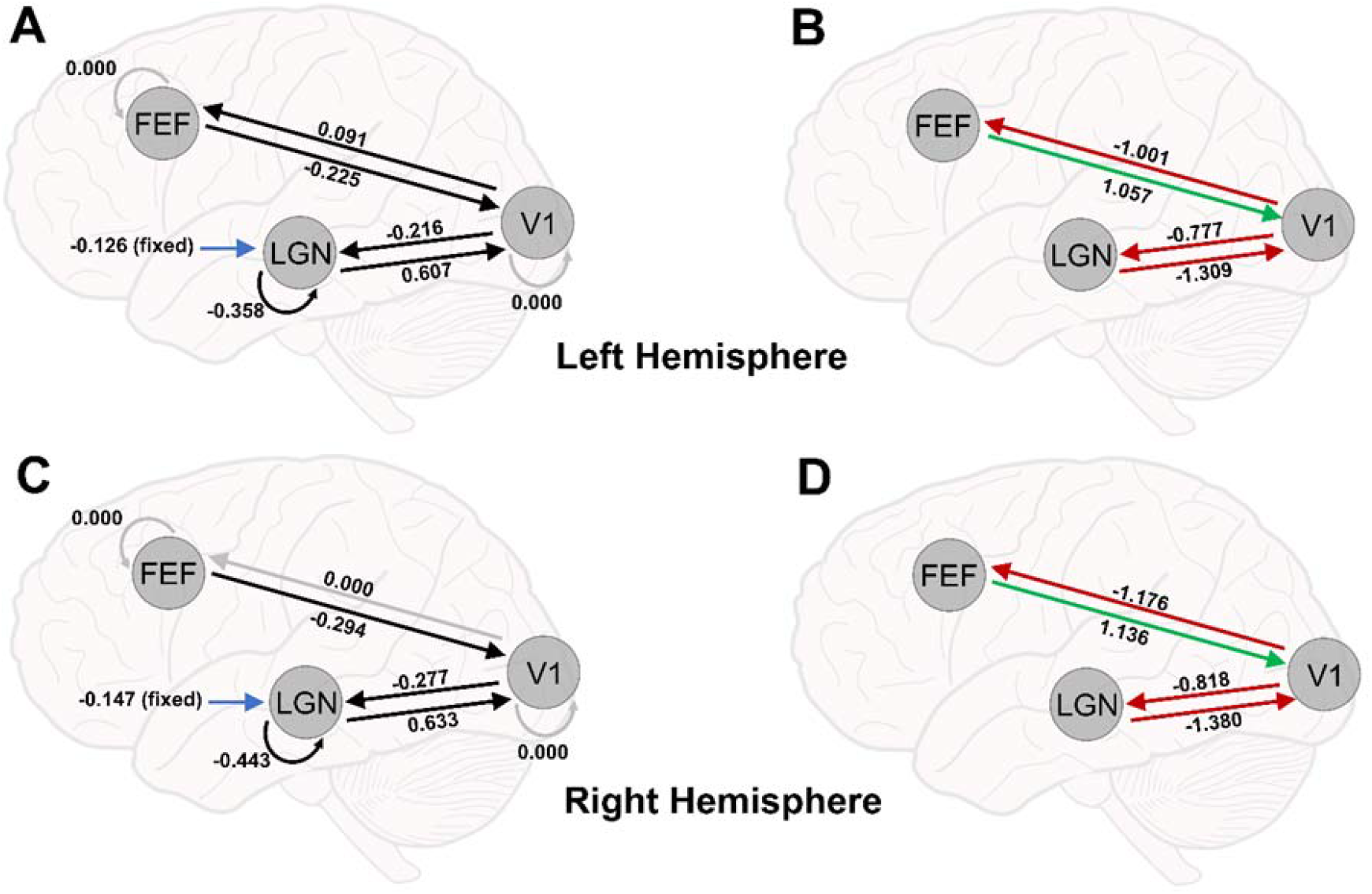
DCM results for LGN – V1 – FEF. (A) Baseline connections for the LGN – V1 – FEF model in the left hemisphere. (B) Modulatory connections for the LGN – V1 – FEF model in the left hemisphere. (C) Baseline connections for the LGN – V1 – FEF model in the right hemisphere. (D) Modulatory connections for the LGN – V1 – FEF model in the right hemisphere. For (A) and (C) solid black arrows with a straight line indicate connections that exceeded posterior probabilities (Pp) of 95%, while grey arrows indicate connections that did not exceed a 95% Pp. Straight arrows represent baseline connections between ROIs, and curved arrows represent the strength of self-connections within ROIs, with smaller (i.e., more negative) values indicating less self-inhibition within the ROI (i.e., long activation sustainment). The significant model input (*C* matrix) is indicated as “free” and “fixed” to LGN. For (B) and (D) red arrows indicate significant negative modulatory effects and green arrows indicate positive modulatory effects (Pp >95%).

## 4. Discussion

Separate lines of evidence have pointed to a role for the MTL memory system as well the oculomotor system in the mental construction of scenes (Mirza et al., 2016; Mullally et al., 2012; Parr & Friston, 2017; Pearson, 2019). Recent computational modeling evidence has shown, using simulated stimulation, that information may rapidly flow from the MTL to regions of the oculomotor system (Ryan et al., 2020a); thereby highlighting an intimate relationship between the two systems (see Ryan et al., 2020b). However, to date, the information flow between the MTL and oculomotor system, and with early visual cortex, for the mental construction of scenes has not been investigated. The current study investigated this effective connectivity using a scene construction task in which participants’ eye movements were either restricted or unrestricted. Since our task did not include external visual input, we were able to determine the directionality of neural connectivity using DCM between key regions of these systems while participants constructed scene representations. As opposed to when eye movements were restricted (*fixed-viewing*), allowing free eye movements (free-viewing) during construction of novel scenes strengthened top-down connections from the MTL to oculomotor regions, and to lower-level cortical visual processing regions, and suppressed bottom-up connections along the visual stream from LGN towards V1 and FEF. Moreover, vividness of imagined scenes was rated higher during free-viewing over fixed-viewing. Taken together, these findings provide novel, non-invasive evidence for the causal architecture between the MTL memory and oculomotor systems associated with constructing vivid mental representations of scenes.

The PPA and HPC have well-established roles in scene processing (Bird et al., 2010; Boccia et al., 2017; Douglas et al., 2017). Given that previous work has demonstrated a positive association between MTL activity and visual exploration (Liu et al., 2017, 2020), and has outlined considerable anatomical connectivity between the MTL memory and oculomotor systems (Ryan et al., 2020a; Shen et al., 2016) we predicted that neural activity in the PPA and HPC would be stronger with unrestricted compared to restricted eye movements. Indeed, both left and right PPA, and left HPC, were more strongly engaged when participants imagined novel scenes in the free-viewing versus fixed-viewing trials. Extensive research has confirmed functional coupling of the PPA and HPC (Baldassano et al., 2013; Sulpizio et al., 2016); here, the current findings highlight the importance of incorporating functional connectivity with oculomotor regions into models of scene construction alongside the MTL. Critically, we revealed a positive modulatory (i.e., excitatory) effect in the free-viewing condition from the MTL towards the FEF. In both hemispheres, the modulatory connections from the HPC and PPA towards FEF were strengthened during free-viewing trials. Likewise, our DCM results revealed that the HPC and PPA received excitatory effects from the FEF when eye movements were unrestricted, which may explain stronger engagement of these MTL regions in the free- viewing condition. Notably, the baseline effective connectivity and the modulation effects from the oculomotor manipulation are largely consistent between the two hemispheres; however, some differences also exist. For example, our results showed positive modulatory (i.e., excitatory) effects unidirectionally towards the FEF from the HPC and PPA in the right and left hemisphere, respectively, but bidirectionally with the FEF in the opposite hemispheres (i.e., left hemisphere for the HPC and right hemisphere for the PPA). Whereas imaging evidence largely suggests hemispheric specialization for spatial contexts in the HPC (Miller et al., 2018; Smith & Milner, 1981) and PPA (Stevens et al., 2012), hemispheric differences in the underlying connectivity of the human MTL with the oculomotor system remain unexplored. Together, these results are highly compatible with the notion that the MTL and oculomotor systems interact in a reciprocal manner (Ryan et al., 2020b), such that information from memory may guide oculomotor behavior (Hannula & Ranganath, 2009; Meister & Buffalo, 2016; Voss et al., 2017), and eye movements may support updating of ongoing scene construction, even in the absence of external visual input (Ringo et al., 1994). Specifically, eye movements may serve to recapitulate the relevant layout and features of an imagined scene brought to mind in MTL memory regions (Ryan et al., 2020b; Wynn et al., 2019).

Similar to the HPC and PPA, the FEF showed stronger activation in the free-viewing versus fixed-viewing condition. Extensive evidence has linked the FEF with the cognitive control of eye movements, in both humans and nonhuman animals (Hanes et al., 1998; Mirpour et al., 2018; Robinson & Fuchs, 1969; Schiller et al., 1979; Selvanayagam et al., 2019). The current results add empirical support to the idea that eye movements may have a critical role in the construction of spatiotemporal content (Hannula et al., 2010), and may promote vivid (re)experiencing (Ryan et al., 2020b; Wynn et al., 2019). Although previous modeling studies described the anatomical connections between the HPC and FEF in primate models (Shen et al., 2016), and simulated information flow between the regions (Ryan et al., 2020a), the current study provides the first functional evidence in humans that MTL regions interact with oculomotor control regions, and specifically, here, to facilitate vivid scene construction. These findings also provide a launching point for future investigations into the critical role of MTL and oculomotor system interactions in service of visuospatial memory and related processes. Depending on the nature of the information required by a specific task (e.g., retrieving or constructing faces, instead of scenes), the interactions between these systems may include different cortical processors alongside the HPC (e.g., the fusiform face area, instead of PPA).

Our DCM results provide novel evidence for the directionality of information flow along the visual stream during scene construction, namely top-down positive (excitatory) modulation from PPA towards V1 when eye movements were unrestricted. Functional connectivity within the scene processing network between PPA and peripheral V1 has been shown to develop in humans as early as 27 days old (Kamps et al., 2020) and may reflect maintained retinotopic organization along the visual stream (Huang & Sereno, 2013). Previous DCM studies have shown strengthened top-down modulatory effects during imagination compared to perception from fronto-parietal regions to early visual regions (Dentico et al., 2014; Dijkstra et al., 2017), but, to our knowledge, no study to date has extended these findings to include MTL memory regions. Moreover, our findings demonstrate that top-down information flow during scene construction may be similar to information flow during mnemonic reconstructive processes (i.e., from HPC towards lower-level visual regions), in reverse of perceptual processes where information flow is thought to originate in early visual regions towards the MTL (Linde-Domingo et al., 2019). Consistent with this interpretation, we found an inhibitory influence in the free-viewing condition from V1 to FEF and PPA/HPC in both hemispheres, reflecting suppression in baseline connections from early visual regions towards the MTL. This suggests that MTL activation is internally initiated and maintained during imagination (Campbell et al., 2018), and inhibition of bottom-up information flow may help to avoid external visual distractions, facilitating scene construction (Benedek et al., 2016; Daselaar et al., 2010).

Although there was limited visual input involved in our task, regions in the early visual pathway (V1 and LGN) showed stronger activation in the free-viewing versus fixed-viewing condition. Although V1 activation in the absence of external input is not an intuitive finding given this region’s role in active vision (Hubel, 1982), it is certainly not a new one (Kosslyn et al., 1995; Miyashita, 1995). In a large-scale meta-analysis of studies involving visual imagery with human participants, Winlove et al. (2018) found consistent activation of FEF and V1; our DCM results demonstrate that this directional relationship is strengthened with unrestricted viewing behaviors. Thus, the FEF, likely in tandem with other oculomotor control regions like the parietal lobe (Rafal, 2006), plays a key role in translating viewing-relevant information of mental representations to eye movement behavior. Additionally, top-down activation from higher-level memory regions to lower-level perceptual regions during mnemonic retrieval has been found in the memory literature (Linde-Domingo et al., 2019; Naya et al., 2001). In rodents, visual regions as early as V1 may process spatial information modulated by the HPC through neural oscillations (Fournier et al., 2020). Our DCM results further suggest that V1 activation can be driven by two top-down pathways: one, along the ventral visual pathway (from PPA to V1) which likely originates from the HPC, and two, through the interaction between the MTL and dorsal oculomotor control system (from HPC through FEF to V1). The specific function of the two pathways should be interrogated in future investigations.

The role of the LGN in unrestricted eye movements was of particular interest in the present study, as there is considerable debate over this region’s role in the construction of scene representations, and how this region may be connected to later visual regions in service of visual imagery (Lesica et al., 2006; Tadmor & Tolhurst, 2000). Previous work suggests that the LGN is primarily related to saccadic control; namely, that saccades in darkness lead to enhanced activity in the LGN, whereas saccades made during strong visual stimulation suppress activity (Sylvester et al., 2005). Similar to modulation effects from V1 to oculomotor control regions, our DCM results showed that the bottom-up influence from LGN towards V1 was inhibited. More importantly, top-down enhancement from the FEF to V1 was not extended to LGN. These results suggest that, in contrast to V1, stronger involvement of LGN in the free-versus fixed-viewing condition was not due to top-down excitations when eye movements were unrestricted. Therefore, different LGN involvement in the two conditions may be primarily related to the eye movement manipulation during the task *per se*, rather than mental construction of scenes. Specifically, the stronger effective connectivity between the LGN and V1, and from V1 to FEF, in the fixed- compared to free-viewing condition is likely driven by the task requirement of maintaining fixation. The bottom-up information flow may help participants to monitor their gaze location and to make adjustments when their gaze deviated from the fixation dot. However, an account based on “effort” is unlikely to fully explain the results here as the relative increase in univariate activity for these regions was observed in the free-viewing condition, rather than the fixed-viewing condition. A “cognitive effort” account would instead predict an increase in BOLD activity in the opposite direction, i.e., greater for fixed- compared to free-viewing.

Likewise, it could be argued that the present results were simply due to instructions that may have led to increased working memory or executive control demands. However, previous work has shown that maintaining central fixation does not appear to significantly increase working memory demands (Armson et al., 2019), perhaps because, even under such instructions, participants still move their eyes, albeit with fewer fixations and smaller saccades, consistent with other work in the literature (Damiano & Walther, 2019; Liu et al., 2020; Welke & Vessel, 2022). Also, here, the fixed-viewing condition did not show obvious differential engagement of the brain regions that support executive control, compared to the free-viewing condition (see Figure 5B). Furthermore, the brain regions used in the DCM analyses each showed stronger activation in the free-viewing than the fixed-viewing condition, rather than the other way around. This result pattern indicates that the engagement of these regions by the free-viewing condition is likely not due to potential differences in working memory demands. Instead, the simplest explanation is that the engagement of these regions is related to oculomotor control during free-viewing, which is fully consistent with, and robustly supported by, the literature (Conti & Irish, 2021; Meister & Buffalo, 2016; Ramkumar et al., 2016; Shen et al., 2016). Moreover, the finding of lower vividness ratings for the mental construction of scenes during fixed-viewing compared to free-viewing are consistent with the neural data showing that the top-down connectivity from HPC/PPA/FEF to lower visual regions was enhanced and the reversed bottom-up connectivity was weakened. Therefore, maintaining fixations likely negatively impacted scene construction due to reduced ability to translate mental representations of novel scenes to viewing-relevant behavior, rather than due to an increased demand on cognitive effort or working memory capacity.

## 5. Conclusions

To summarize, in the present study, we successfully applied DCM to investigate the causal interactions between the MTL memory and oculomotor systems in support of scene construction. Our findings provide strong support for a top-down influence from the MTL to oculomotor control region FEF and to early cortical, but not subcortical, visual regions, and an inhibitory bottom-up modulatory effect of visual exploration from LGN to V1 and FEF when a mental scene representation was constructed. More generally, this work demonstrates how the MTL may guide eye movements to support vivid, experiential phenomena during imagination and recollection. Eye movements, as such, may be a natural effector system for memory (Ryan & Shen, 2020), and critical for the mental imagery of scenes.

## Acknowledgements

We would like to thank Ryan Aloysius, Arber Kacollja, Ling Li, and Mandy Ding for their help at different stages of this research project.

## CRediT Author Contributions

Natalia Ladyka-Wojcik: Conceptualization; Data curation; Formal analysis; Investigation; Methodology; Validation; Visualization; Writing – Original draft; Writing – Review & editing Zhong-Xu Liu: Conceptualization; Data curation; Formal analysis; Investigation; Supervision; Methodology; Validation; Visualization; Writing – Original draft; Writing – Review & editing Jennifer D. Ryan: Conceptualization; Funding acquisition; Project administration; Resources; Supervision; Writing – Review & editing

## Data availability statement

Analysis scripts and final results matrices are openly available on Open Science Framework (OSF) at https://osf.io/nt45v/ (doi: 10.17605/OSF.IO/NT45V). All processed data associated with reported results are available on request from the corresponding authors, NLW or ZXL. The original data are not publicly available due to ethics restrictions at the time of data collection.

